# An outbreak of SARS-CoV-2 in big hairy armadillos (*Chaetophractus villosus*) associated with Gamma variant in Argentina three months after being undetectable in humans

**DOI:** 10.1101/2022.08.23.503528

**Authors:** Franco Lucero Arteaga, Mercedes Nabaes Jodar, Mariela Mondino, Ana Portu, Monica Boeris, Ana Joly, Ana Jar, Silvia Mundo, Eliana Castro, Diego Alvarez, Carolina Torres, Mariana Viegas, Ana Bratanich

## Abstract

The present pandemic produced by SARS-CoV-2 and its variants represents an example of the one health concept in which humans and animals are components of the same epidemiologic chain. Animal reservoirs of these viruses are thus the focus of surveillance programs to monitor their circulation and evolution in potentially new hosts and reservoirs. In this work, we report the detection of SARS-CoV-2 Gamma variant infection in four specimens of *Chaetophractus villosus* (big hairy armadillo/armadillo peludo) in Argentina. In addition to the finding of a new wildlife species susceptible to SARS-CoV-2 infection, the identification of the Gamma variant three months after its last detection in humans is a noteworthy result, raising the question of potential unidentified viral reservoirs.

## Introduction

The SARS-CoV-2 pandemic has triggered investigations on the potential role of animals, and in particular, wildlife, could play as reservoirs of this virus since they may facilitate the generation of new variants with novel capacities as human pathogens (1,2). Thus, constant monitoring of circulating SARS-CoV-2 variants in any animal species, including humans is essential to detect those that may have changed their host spectrum compared to the index virus (genomic sequence of SARS-CoV-2 identified from the first human cases, December 2019) (https://www.who.int/en/activities/tracking-SARS-CoV-2-variants).

The World Health Organization has classified some groups of SARS-CoV-2 sequences as variants of concern (VOC), naming them as Alpha, Beta, Gamma, Delta and Omicron (https://www.who.int/en/activities/tracking-SARS-CoV-2-variants). Even though, according to the level of circulation and reporting to international public databases, at beginning of March 2022 they were also defined as “Currently circulating VOCs” (Omicron), and “Previously circulating VOCs” (Alpha, Beta, Gamma, Delta) (https://www.who.int/en/activities/tracking-SARS-CoV-2-variants).

Worldwide, VOC Gamma has been reported from April 2020 to January 2022, and in Argentina, from July 2020 to December 2021. On the other hand, Gamma sequences have been classified as lineage P.1 and several derivatives lineages under the Pango system (https://www.who.int/en/activities/tracking-SARS-CoV-2-variants).

In Argentina, since March 2020, there has been an integrated health surveillance system (SNVS), where the number of positive cases of COVID-19, hospitalizations and deaths are registered. It also provides information on the molecular characterization of the variants that circulate in the country and are detected by partial and complete genomic sequencing and specific RT-qPCR techniques for VOCs and variants of interest (VOIs). In addition, PAIS is the inter-institutional federal consortium of SARS-CoV-2 genomics in Argentina, which was created by the Ministry of Science and Technology to monitor SARS-CoV-2 diversity and evolution in the country, including surveillance of SARS-CoV-2 variants of public health interest and notification to the SNVS (http://pais.qb.fcen.uba.ar/). Based on this information, the Argentine Ministry of Health regularly publishes reports informing the situation of the circulation of the variants in the country.

According to the OIE latest report SARS-CoV-2 has been detected in twenty-three animal species: cats, dogs, mink, otter, pet ferrets, lions, tigers, pumas, snow leopards, gorillas, white-tailed deer, fishing cats, Binturong, South American coati, spotted hyena, Eurasian lynx, Canada lynx, hippopotamus, hamster, mule deer, giant anteater, West Indian manatee, and black-tailed marmoset (https://www.who.int/publications/m/item/scientific-advisory-group-on-the-origins-of-novel-pathogens-report). In some of these cases, the SARS-CoV-2 variant affecting these species has been identified (Scientific Advisory Group for the Origins of Novel Pathogens (SAGO) report June 2022) (https://www.who.int/publications/m/item/scientific-advisory-group-on-the-origins-of-novel-pathogens-report). In Argentina, infections of cats, dogs and pumas with SARS-CoV-2 have been reported and at least in the feline case, the lineage identified in the animal by the PAIS consortium was in agreement with one of those circulating in humans at the same moment (B.1.499 lineage) (4).

In this work we report for the first time, the infection with SARS-CoV-2 of big hairy armadillos/armadillo peludo (*Chaetophractus villosus)* during the third wave of COVID-19 in Argentina (March 2022). This species inhabits the Pampas region, located in the centre of Argentina, and characterized by wide plains dedicated largely to livestock and agricultural production. The advance in production systems on the natural ecosystems of the area have favoured the coexistence of wildlife, domestic fauna, and human beings. Different species of rodents and birds, wild pigs, wild cats and armadillos, among others, are part of the wild diversity of the region. Among the armadillos that inhabit this area, the *Chaetophractus villosus* stands out; it is commonly known by the name of “peludo pampas” or “big hairy armadillo” and belongs to the mammalian Cingulata order. It has a wide distribution throughout the country, generally limited to arid and semi-arid regions that range from southern Bolivia and Paraguay to the Isla Grande of Tierra del Fuego in Argentina. However, their remarkable ability to adapt to different climates and diverse food resources has allowed them to establish themselves in more humid places such as the agricultural region of the Pampas plains, which has meant a problem for crop production in the area.

We also report in this work that, surprisingly, these *Chaetophractus villosus* specimens were infected with the SARS-CoV-2 Gamma variant, a variant associated with the second wave of COVID-19 in Argentina (5) and a long time after its replacement by Delta and Omicron variants in humans. In the last report published by the Argentine Ministry of Health in April 2022, it is informed that the Gamma variant stopped being detected in the population around epidemiological week 51 of 2021 with 0.1% of the total cases characterized up to that moment (April 2022).

The findings described here have important implications for the epidemiology of this viral infection and the role wildlife may play as the source of potential reservoirs.

## MATERIALS AND METHODS

### Animal samples collection and processing

Four six months old *Chaetophractus villosus* specimens (A#1, A#2, A#3 and A#4) were housed in a building of the Faculty of Veterinary Sciences of the University of La Pampa (La Pampa province, Argentina); three of the animals had been donated by rural workers in the area, and the fourth one was born in the facilities. The four animals had been at the university facilities since April 2021 and were being bred as part of a collateral project related to brain neurochemistry. At the time, oropharyngeal/nasal/rectal swabs and faecal samples were also obtained for a surveillance project to study potential novel coronaviruses in wildlife (with negative results).

University personnel took care of the animals, which were maintained with balanced feed and drinking water ad libitum. All four animals were housed in the same enclosure, with a heated environment and a chip bed. The building was undergoing renovations and due to the pandemic, the construction work had been momentarily stopped.

The animals started with respiratory symptoms during the second week of March 2022, with signs compatible with rhinitis; there was nasal discharge and sneezing in three of the animals (A#1, A#2 and A#4), and only one of them had diarrhoea (A#4). There was no evidence of increased body temperature. At the onset of symptoms, the animals also experienced a heavy parasitic infestation that was later diagnosed as gastrointestinal nematodes. During the symptomatic period beginning on March 14, 2022, oropharyngeal swabs were taken twice (March 14 and 29) and rectal swabs were obtained four times (March 19, 28 and 29, and April 1) (Fig.1). At the time of the onset of symptoms, none of the staff members handling these animals was tested for the presence of SARS-CoV-2 neither they showed respiratory symptoms but all of them had been vaccinated twice. All samples were kept in 500 ul of DNA/RNA Shield 1x reagent (Zymo Research) until processed.

In addition, serum samples were taken 15 (all 4 animals) and 30 days (A#1 and A#4) after the onset of symptoms to evaluate the presence of total antibodies and their neutralizing activities. For the extraction of blood samples, each animal was anesthetized following an anaesthetic protocol of Xylazine, Ketamine and Diazepam by intramuscular injection. Blood extraction was performed by catheterization of the saphenous vein. After the anaesthetic procedures for the sera collection, two of the animals (A#2 and A#3) perished. A necropsy was performed, and liver, spleen and lung tissue samples were taken from both animals for RNA extraction with TRizol® Reagent (Thermo Fisher). RNA was used for RT-qPCR assays as described below.

All animal studies and procedures were performed according to an approved protocol by the University of La Pampa, CAICUAE (Internal Advisory Commission for the Care and Use of Experimental Animals), University of La Pampa, Argentina.

### RT-PCR/ RT-qPCR assays on animal samples

For oropharyngeal and rectal swabs, 100 ul were used for RNA extraction (Quick-RNA Viral Kit, Zymo Research) whereas for faeces, a dilution 1/10 in DNA/RNA Shield 1x reagent was previously done. cDNA synthesis was performed with a RT kit (EasyScript First-Strand cDNA Synthesis Super Mix, Transgene Biotech) with random hexamers according to the manufacturer’s instructions. cDNA was utilized for a pan-corona PCR that targets a conserved RdRp region of approximately 440-bp (6). The nested PCR (nPCR) was performed with 5 ul of cDNA, 200 uM dNTPs, 1x GoTaq buffer, 1 unit of GoTaq enzyme (Promega) and 25 pmoles of pan-coronavirus primers (10). In a first round, primers RdRp For1 5’GGKTGG GAYTAYCCKAARTG 3’ and RdRp Rev2 5’TGYTGTS WRCARAAYTCRTG 3’ were used. For the second round, 5 ul of the first round were utilized with primers RdRp For3 5’ GGTTGGGACTATCCTAAGTGTGA 3’ and RdRp Rev4A 5’ CCATCATCAGATAGAATCATCAT 3s’. Both PCR reactions were run with the following cycling conditions: 2 minutes at 94°C, 40 cycles of 94°C 1 minute, 48°C (first round) or 55°C (second round), 1 minute 72°C and a final cycle at 72°C 5 minutes. PCR products were purified and submitted for sequencing by the Sanger method (INTA Castelar, Argentina)

All samples (negative and positive) were re-tested with a RT-qPCR targeting a 127-bp region of the SARS-CoV-2 E gene. RT-qPCR was performed with 5uL of RNA extract and 15uL of supermix (SuperScript™ III Platinum™ One-Step RT-qPCR Kit, Invitrogen™), using primers for E_Sarbeco_F 5‘ACAGGTACGTTAATAGTTAATAGCGT 3’ and Rev E_Sarbeco_R 5’ ATATTGCAGCAGTACGCACACA 3’ (7).

### Sera collection, ELISA and neutralization assays

SARS-CoV-2 IgG antibodies were tested by an indirect ELISA using pCDNA3-SARS Clone 1 S-RBD expressed in HEK-293 cells. This clone spans the spike RBD region (amino acids 319 to 541) of a local SARS-CoV-2 positive sample (B1 variant) fused to the N-terminal S protein signal sequence. In the assay, antibodies were detected using Thermo Scientific™ Pierce™ Recombinant Protein A, Peroxidase at a 1:25000 dilution. Optical density (OD) was measured at 450/630 nm and EU were calculated as positivity percentage relative to human positive and negative controls (gently provided by Dr. Andrea Gamarnik, Instituto Leloir, Buenos Aires, Argentina).

For the neutralization assays Vero E6 cells were cultured at 37°C and 5% CO2 in complete Dulbecco’s Modified Eagle’s Medium (DMEM, Gibco) containing 10% (vol/vol) foetal bovine serum (FBS, Internegocios, Buenos Aires, Argentina), 100 lU/mL penicillin, and 100 μg/mL streptomycin (Gibco) (DMEM10). SARS-CoV-2 SARS-CoV-2 D6124G variant (B.1 lineage) and the Gamma variant (P.1) used in the assays were isolated from nasopharyngeal specimens at the Instituto de Investigaciones Biotecnológicas, UNSAM, CONICET and adapted in Vero E6 cultures. Cells were seeded in 96-well plates at a density of 1,5 × 104 cells per well in complete media and incubated 24h at 37°C and 5% CO2. SARS-CoV-2 (300 TCID50) were preincubated with serially diluted sample sera for 1h at 37°C starting at a 1:4 sample dilution. Each sample dilution was tested by duplicate. Then, virus-sera mixture was added onto Vero E6 cells in a final volume of 100 µl in DMEM 2% FBS. Infected cells (virus control, VC) and mock infected cells (cell control, CC) were added as controls. After a 72h incubation at 37°C and 5% CO2, cultures were fixed with formaldehyde 3% at 4°C for 24h and stained with crystal violet solution in ethanol. A surface scan at 595 nm was performed in each well in a microplate reader (FilterMax F5, Molecular Devices, California, USA). Average optical density (OD) of each well was used for the calculation of the percentage of neutralization of viral CPE for each sample (S) as: Neutralization% = 100 x (DO sample - DO virus control) / (DO cell control - DO virus control).

### Whole genome sequencing of SARS-CoV-2 positive samples from *Chaetophractus villosus* specimens

The complete genome sequencing of SARS-CoV-2 from *Chaetophractus villosus* positive samples was performed with the Midnight RT-PCR protocol, for use in Oxford Nanopore MinION (PCR tiling of SARS-CoV-2 virus with rapid barcoding and Midnight RT PCR Expansion -SQK-RBK110.96 and EXP-MRT001): https://community.nanoporetech.com/protocols/pcr-tiling-of-sars-cov-2-virus-with-rapid-barcoding-and-midnight. Sequences were submitted to GISAID: hCov-19/ *Chaetophractus villosus/*Argentina/PAIS-A1415/2022I2022-03-29 l La Pampa (A#1) (EPI_ISL_14425591); hCov-19/ *Chaetophractus villosus*/Argentina/PAIS-A1416/2022 I 2022-03-29 l La Pampa (A#2) (EPI_ISL_14425592), and hCov-19/ *Chaetophractus villosus*/Argentina/PAIS-A1417/2022I2022-03-29 l La Pampa (A#4) (EPI_ISL_14425830)

### RT-qPCR and sequencing on SARS-CoV-2 positive human samples

A retrospective analysis of human positive cases of COVID-19 was carried out between 2/15/2022 and 4/28/2022 in the city of General Pico and surroundings where the animals were located (La Pampa province, Argentina). A total of 36 RNA samples extracted from human nasopharyngeal swabs were recovered from a collection of positive samples from the province, stored at -70°C at the laboratory of the Department of Epidemiology of La Pampa.

A variant-specific RT-qPCR with the TaqMan™ SARS-CoV-2 kit Mutation Panel (Applied Biosystems) was used to screen the VOC/VOI mutations S_L452R and S_T478K following manufacturer ‘s protocol.

For partial sequencing of the gene that encodes for the Spike protein, the traditional Sanger method was used, with the sequencing protocol recommended by the CDC (https://wwwnc.cdc.gov/eid/article/26/10/20-1800_article), which amplifies segment 29 of the mentioned protocol (fragment comprised between amino acids S_428 and S_750).

### Phylogenetic analysis

An initial assignment of Pango lineages on the three sequences was performed using the Pangolin COVID-19 Lineage Assigner tool (https://pangolin.cog-uk.io/; Pangolin v4.0.6, pango-designation v1.10), resulting in lineage P.1.15 (VOC Gamma).

Thus, a phylogenetic analysis was performed on a dataset of SARS-CoV-2 sequences that included: reference sequences of the main Pango lineages, focusing on P.1 and its derivatives, the three sequences of interest, and all the sequences retrieved from Audacity Instant application in GISAID (http://www.gisaid.org) on to June 20, 2022) (n=128), which searches the entire EpiCoV database to find closely related sequences (those with the lowest number of SNPs).

The alignment was built with MAFFT v7.486 (8) inspected and edited with BioEdit (9), and the maximum likelihood tree was built with IQ-TREE v.2.1 (10). The molecular evolution model appropriate for the dataset was estimated with ModelFinder (11) and the reliability of branches and clusters was evaluated using the SH-approximate likelihood ratio test (1000 replicates) and Ultrafast bootstrap Approximation (UFB) (10000 replicates) (12,13).

## Results

### PCR and serology results of animal samples

Three of the oropharyngeal samples preliminary obtained on 03/14/2022 (A#1, A#2 and A#4) were positive by the pan-corona RT-PCR assay. Sequencing of the PCR products confirmed the presence of SARS-CoV-2. Pooled individual rectal swabs (from 03/28/2022, 03/29/2022 and 04/01/2022) and oropharyngeal swabs (03/29/2022) were tested in a second stage. Most samples resulted positive by the pan-corona PCR assay and the RT-qPCR except for the samples from A#3 (Fig 1). RT-qPCR Ct values of positive oropharyngeal/rectal samples were between 22.1 and 24.

**Figure 1.**
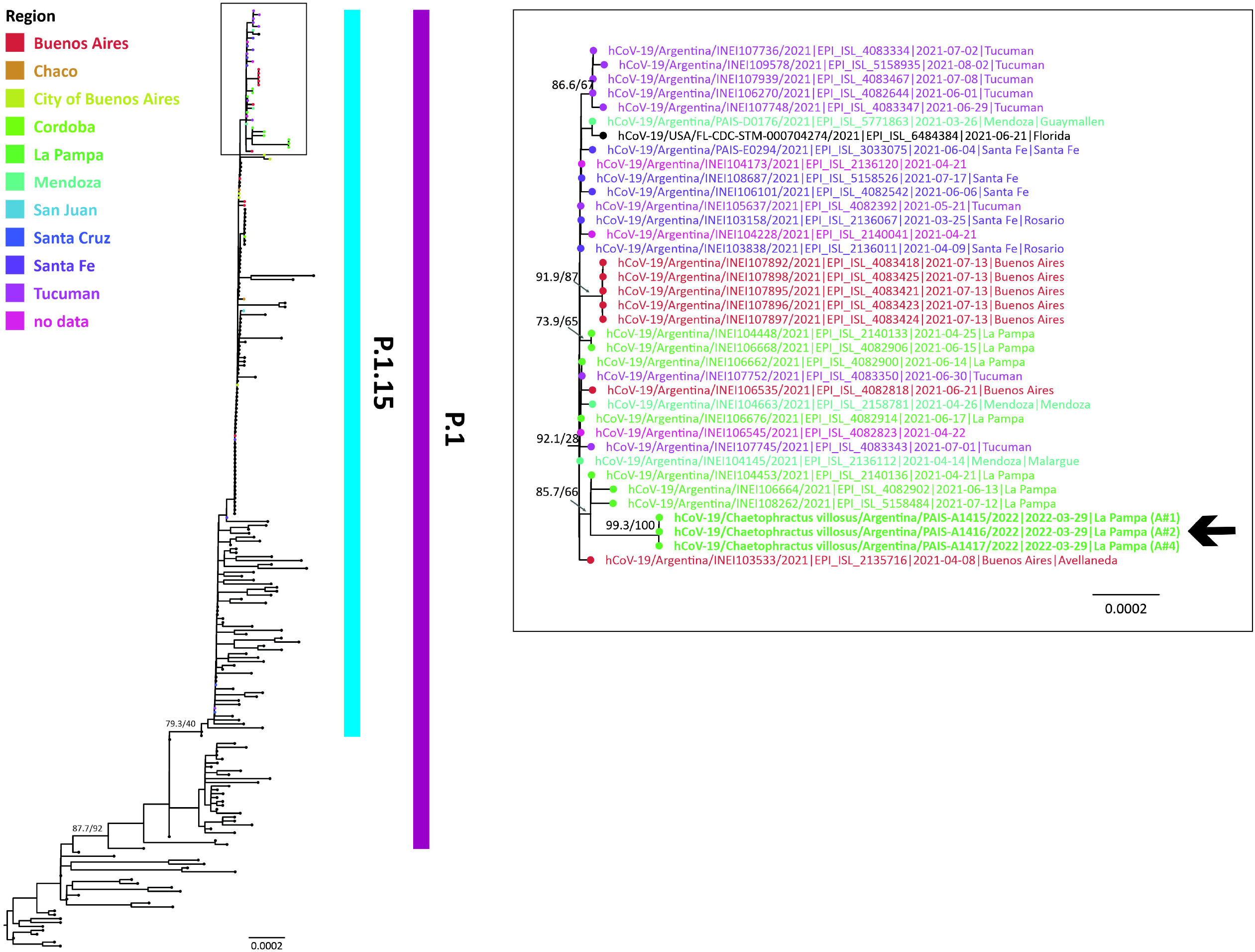
Timeline of SARS-Cov-2 infection in *Chaetophractus villosus*, initiation of symptoms, collection of samples and techniques utilized with their results.

Necropsy of the deceased animals did not show any macroscopic lesion, particularly in A#2 which was the one with respiratory symptoms (A#3 was always asymptomatic). Lung, spleen, and liver tissues from this animal were tested by RT-qPCR. Interestingly, the spleen was positive whereas the lung gave inconclusive results (Cts >40). The liver sample was negative.

Serum samples from A#1, A#2, A#3, and A#4 obtained on 03 /29/2022 gave positive reactivity by the SARS-CoV-2 RBD ELISA. An increment of reactivity was also detected in A#1 and A#4 two weeks later.SARS-CoV-2 specific neutralizing antibodies were detected at low levels 15 days post-onset of symptoms in all animals. The serum samples from the two remaining animals (A#1 and A4) also showed neutralizing activity, in agreement with the ELISA and the PCR results. A#3, although showing a positive ELISA and some neutralizing activities, was always negative by all PCR tests (swabs and tissues) (Fig 1) (Table 1).

### SARS-CoV-2 whole genome sequences

Complete or nearly complete genome sequences of three animals (A#1, A#2 and A#4) were obtained. The samples sequenced were a rectal swab pool from A#1 with a Ct value of 24.35, and oropharyngeal swabs with Ct values of 22.3 and 22.1 from A#2 and A#4 respectively.

Compared to the Wuhan-Hu-1/2019 reference sequence (EPI_ISL_402124), the three sequences showed 42 nucleotide substitutions in different viral genes: 18 in ORF1a/1b, 12 in S, 8 in N, 1 in ORF13a, 1 in ORF6, 1 in ORF8 and 1 in 5’UTR, resulting in 25 amino acid replacements as indicated in Table 2. Of these, 21 amino acid changes corresponded to the constellation of mutations designated by the WHO as characteristic of the Gamma variant (https://www.who.int/en/activities/tracking-SARS-CoV-2-variants/).

**Table 2.** Location of nucleotide differences found in the SARS-COV-2 genome sequences from Armadillos compared to the reference sequence of hCoV-19/Wuhan/WIV04/2019 (EPI_ISL_402124). Nucleotide changes at the analysed positions of the 3 most related sequences identified by phylogeny are also shown. Gray shaded text indicates genome positions with nucleotide changes that are unique to the Armadillos genome sequences. Bold text indicated the profile of amino acid changes designated by WHO as characteristic of the gamma variant (lineage P.1). NA, not applicable, as nt change in the noncoding region; ND, not determined; ORF, open reading frame; NSP, non-structural protein.

| Date of sampling 29/3/22 |  |  |  | 12/4/22 |  |  |
| --- | --- | --- | --- | --- | --- | --- |
| Animal | RBD-ELISA<br>EU | sVN<br>%<br>WT | sVN<br>%<br>Gamma | RBD-ELISA<br>EU | sVN<br>%<br>WT | sVN<br>%<br>Gamma |
| #1 | 54,9 ± 5,8 | 64.2 | 51.6 | 37,3 ± 3,4 | 34.8 | NE |
| #2 | 163,0 ± 9,5 | 11.1 | 25.2 | NE | NE | NE |
| #3 | 54,8 ± 14,7 | 3.2 | 25.9 | NE | NE | NE |
| #4 | 38,9 ± 4,4 | 22.7 | 69.3 | 53,2 ± 10,3 | 50.0 | NE |

Remarkably, compared to the most related genome sequences of the Gamma variant, the three genome sequences of SARS-CoV-2 of this study displayed six single-nucleotide polymorphisms (SNP) distributed in 3 genes, the gene coding for replicase polyprotein (ORF1ab), nucleocapsid phosphoprotein (N) and ORF6. Overall, ORF1ab harboured three synonymous mutations, located in NSP2, NSP3, NSP13 and one non-synonymous mutation (NSP12_R905K). Likewise, the nucleocapsid phosphoprotein and ORF6 harboured two non-synonymous mutations (N_S194L and ORF6_I14T) (Table 2).

### Phylogenetic analysis

A phylogeny tree was built to analyse the evolutionary relationship of the SARS-CoV-2 sequences from the three *Chaetophractus villosus* specimens to the most related sequences obtained from GISAID. The three sequences belonged to the Gamma variant, lineage P.1.15, and formed a highly supported monophyletic group compatible with a common source of infection and a unique chain of transmission (Figure 2). The most related sequences to this group correspondeded to samples from the province of La Pampa, Argentina, obtained from human clinical samples between April-July 2021. All the sequences from La Pampa, Argentina, shared a common ancestor with sequences from other regions of the country (Figure 2).

**Figure 2.**
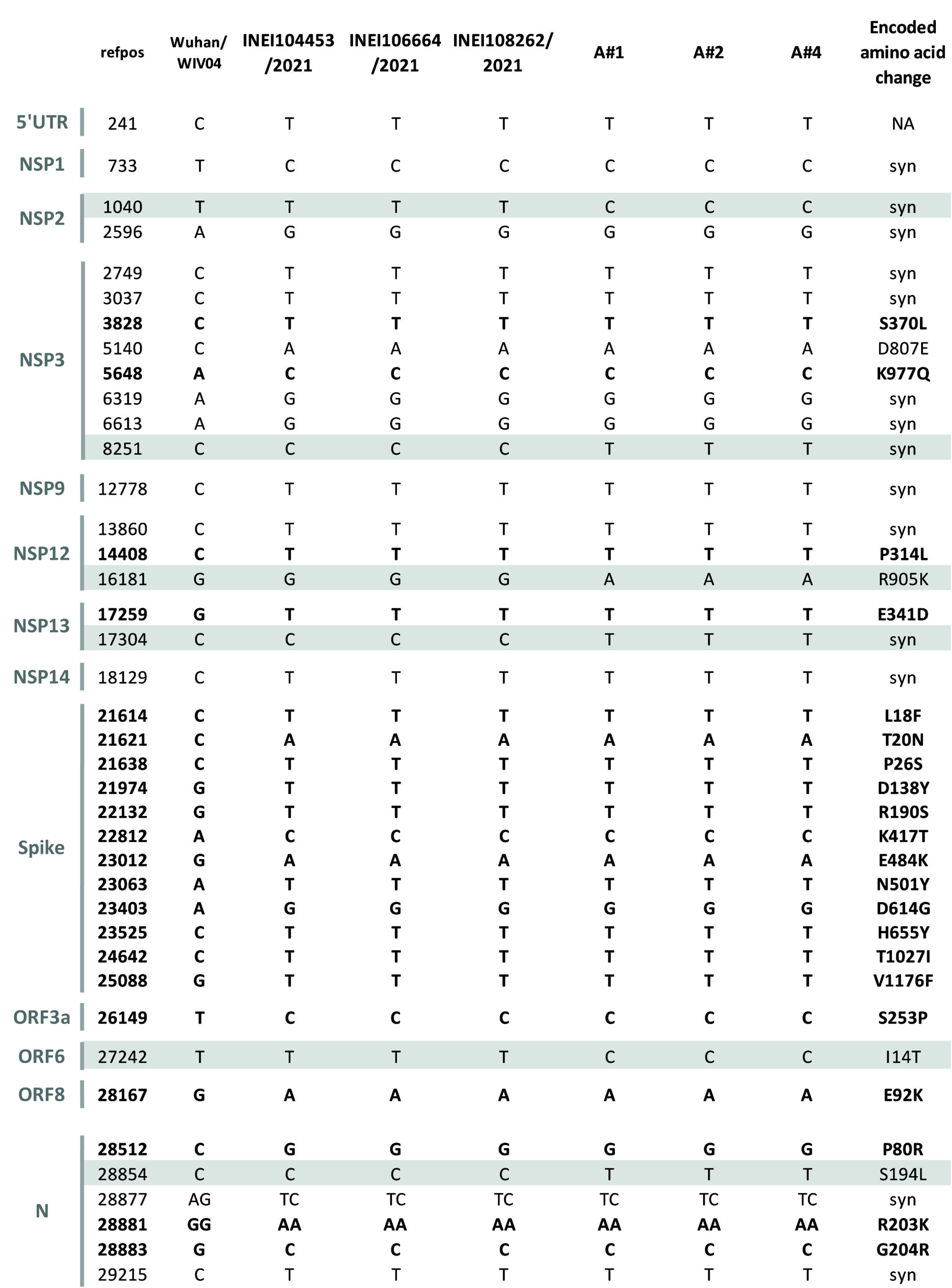
Phylogenetic tree of SARS-CoV-2 whole-genome sequences of Gamma variant (lineage P.1 and derivatives). The largest group with Argentinean sequences is shown. The SH-aLRT/UFB values for the relevant groups are indicated. Sequences from three animals analysed in this work are shown in bold and indicated with a black arrow (A#1, A#2, and A#4).

### Human samples

Due to the fact that the characterization of the variants is not carried out in 100% of the cases of COVID-19, and with the aim of being able to detect if the Gamma variant was still circulating in humans at a very low rate at the time it was detected in these animals, additional human SARS-CoV-2 positive samples from the same region were analysed by variant-specific RT-qPCR and sequencing of the spike protein gene.

Thirty-one of 36 samples had undergone the variant-specific RT-qPCR and all were compatible with the Omicron variant. Subsequently, 16 of them that had RNA with enough quantity and quality were sequenced. In all the sequenced samples, the constellation of mutations detected indicated that all of them corresponded to the Omicron variant, lineage BA.1, showing that there were not cases of Gamma variant during the period, at least in the analysed samples.

## Discussion

We have described the infection of four *Chaetophractus villosus* specimens in March 2022 with the Gamma variant (lineage P.1.15) of SARS-CoV-2 in Argentina. The findings reported in this communication are remarkable from several points of view. The molecular detection confirmed in some tissues (spleen of A#2) and oropharyngeal and rectal swabs in A#1, A#2 and A#4, together with the symptomatic infections and the presence of ELISA specific and neutralizing antibodies in all animals, demonstrate that they had been actively infected with SARS-CoV-2.

To our knowledge, there are no published reports on *Chaetophractus villosus* or other members of the Cingulata order that have been tested for the presence of SARS-CoV-2, antibodies or have been used for experimental infections. Studies on the potential susceptibility of different animal species based on their ACE2 sequence have reported a potential very low susceptibility of the nine banded armadillo (*Dasypus novemcinctus*), a relative of the big hairy armadillos although they belong to different families (14). We believe that the extent and the speed in which this infection progressed among these animals (low Ct values and presence of specific antibodies) suggest that this species is highly susceptible. The identification and sequencing of the *Chaetophractus villosus* ACE2 is in progress to answer these questions.

The other and perhaps, most astonishing result is the finding that these animals were infected with a SARS-CoV-2 Gamma variant, which according to GISAID reports, registered its last record of circulation by January 10, 2022, in Peru (EPI_ISL_9769405) and by December 22, 2021, in Argentina (EPI_ISL_9192218). Furthermore, reports of the PAIS project and the Ministry of Health on the surveillance of SARS-COV-2 by partial sequencing and variant-specific RT-qPCR showed that there were no circulation records of the Gamma variant on those dates. To further investigate a link to circulation of the Gamma variant, a collection of human positive samples was obtained from the city and surroundings in where the animals were located (General Pico, La Pampa, Argentina) in the same period to investigate whether cryptic circulation of this variant was happening without detection, resulting all samples as Omicron variant. The results would suggest that the personnel or other individual handing these animals seem not to have been the most probable source of infection although indirect contact through fomites cannot be discarded as has been mentioned in the case of positive deer in USA (15). Other animals that could fortuitously enter the facility might have initiated the outbreak. Personnel suggested that among them, rodents would be the most probable ones since they could have transited undetected through the facility and even be ingested by these animals. Since in some opportunities, faeces were observed close to where the food stocks were kept, this observation would support the rodent hypothesis. Pigeons have also been mentioned as possible visitors although avian species in general are considered as not susceptible to SARS-CoV-2 infection (16, https://www.who.int/publications/m/item/scientific-advisory-group-on-the-origins-of-novel-pathogens-report). Whether these observations play a role in the described *Chaetophractus villosus* incident requires further investigation. Transmission among different animal species has been previously reported (17).

Even though the uncertainty on the source of infection, the sequencing data from the three animals clearly points to a single transmission event to the animals with low intra-cluster diversity. The non-synonymous mutations identified in *Chaetophractus villosus* have not been reported previously in other animals infected naturally or experimentally with SARS-CoV-2 Gamma variant. A change in N 194 has been described in human sequences as N_S194L (with a potential association with symptomatic patients (18) but in experimental infections in cats, dogs and mice the change is different (N_S194T) (19). There are no records of ORF6_ I14T and NSP12_ R905K changes in SARS-CoV-2 infected animals. The impact of these aminoacidic differences is thus, unknown at this stage.

The discovery of *Chaetophractus villosus* as a susceptible host of SARS-CoV-2 infection has enormous relevance due to the increasing transit of this species from the wild to populated areas. *Chaetophractus villosus* has great plasticity in terms of its eating habits; they are omnivorous animals par excellence. Their diet is based on the consumption of products of plant origin (reason for which they often destroy grain storages in livestock farming systems) but they also can consume small animals of various types such as rodents, reptiles, and birds. Likewise, it has been proven that *Chaetophractus villosus* has scavenging habits, consuming the carcasses of different animals, aborted foetuses, remains of the placenta, etc. Specimens of this species have also been found in local cemeteries, where they have caused the destruction of coffins of deceased people, which suggests that human corpses can also be a food source for *Chaetophractus villosus*. In addition to this, these animals are consumed in an artisanal way by local people throughout the country, which could suggest an important role in the permanence and dissemination of different pathogens. The finding described in this work suggests that one of those pathogens could be SARS-CoV-2 since this species could be infected by the Gamma variant and be transmitted efficiently among its members. We do not know if these animals can transmit the infection to humans, but with this work we were able to determine that *Chaetophractus villosus* or other animals not yet identified could act as reservoirs of SARS-CoV-2 variants.

## Acknowledgments

We would like to thank doctors Claudia Rechimont, José Carlos Usero, Matías Villasana and María de los Ángeles Irastorza from the laboratory of the Department of Epidemiology (La Pampa province) for the data from the surveillance of variants by RT-qPCR and for sending human samples for sequencing. We gratefully acknowledge the Authors from the Originating laboratories responsible for obtaining the specimens and the Submitting laboratories where genetic sequence data were generated and shared via the GISAID Initiative, on which part of this research is based.

## Author contributions

FLA and AB prepared the draft manuscript. FLA, MM, MP and MB oversaw the collection of samples. FLA and AB carried on the processing of samples and the first part of the viral identification. AJ (Ana Joly) AJ (Ana Jar) and SM produced the RBD protein and performed the ELISA assays.EC y DA performed the seroneutralization assays. MNJ and MV performed the whole genome sequencing of the positive swabs and variant determination of human samples from the La Pampa province. CT did the phylogenetic analysis. MNJ, MV and CT revised the manuscript. The final version of the manuscript was approved by all authors.

## Data availability

A list of accession numbers of the sequences used, with information of the contributions of the submitting and the originating laboratories, can be retrieved through the Data Acknowledgement Locator at https://www.gisaid.org with ID EPI_SET_20220627we).

## Figure captions

Table 1. Antibody response obtained with *Chaetophractus villosus* serum samples by RBD-ELISA and seroneutralization. ELISA Units (EU) were calculated as positivity percentage related to human positive and negative controls. Average DO of each well was used for calculation of the percentage of neutralization of viral CPE for each sera dilution (S) as: Neutralization% = 100 x (DO S - DO CV) / (DO CC - DO CV). Sera were tested at 1:8 dilution against WT and at 1:4 dilution against Gamma SARS-CoV-2. NE: not evaluated (no sample).

---

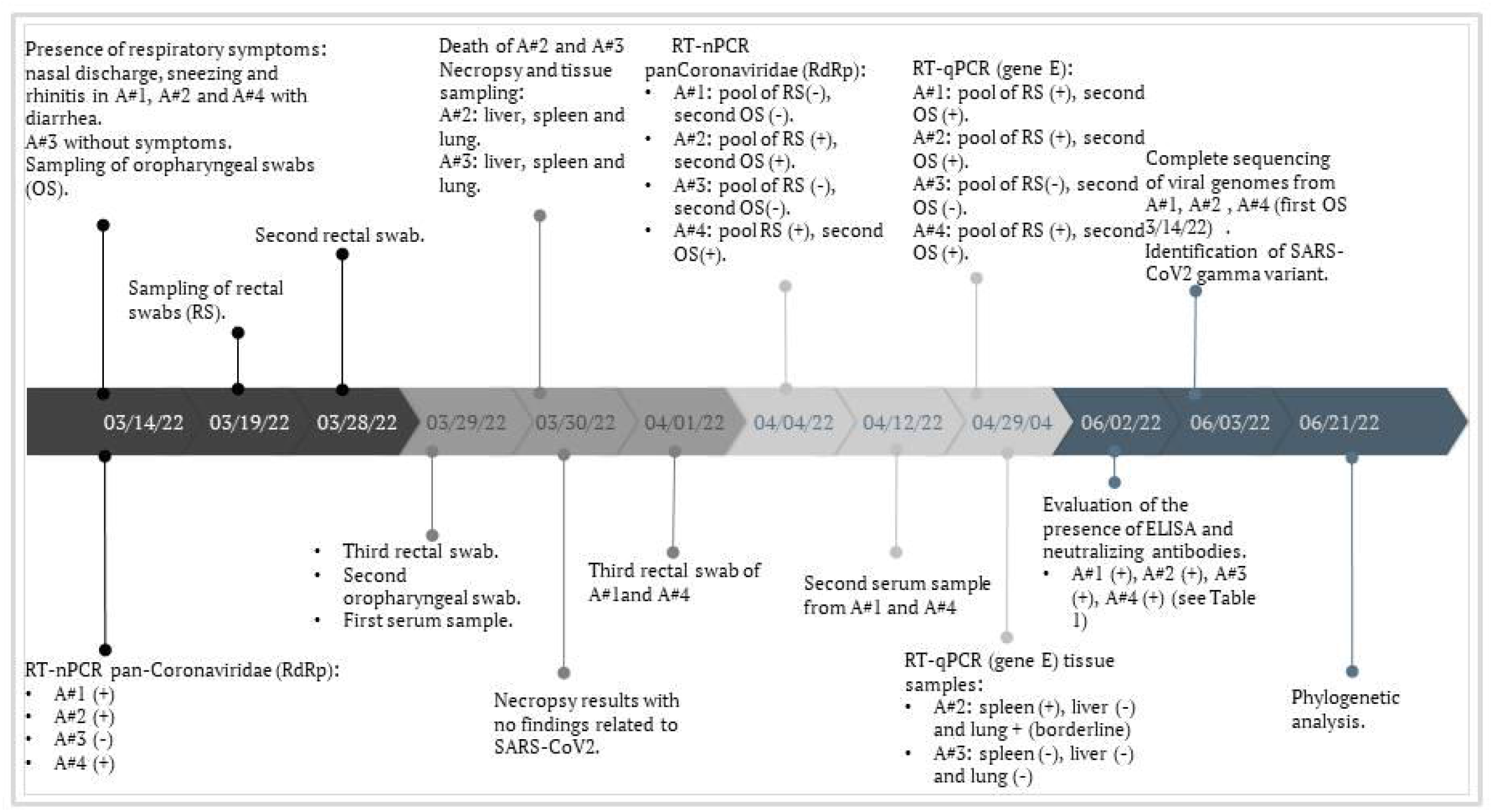

